# Hyper-Cryptic Radiation of a Tropical Montane Plant Lineage

**DOI:** 10.1101/2023.04.26.538369

**Authors:** Ingrid Olivares, Sergio Tusso, MarÍa JosÉ SanÍn, Michael Kessler, Kentaro K. Shimizu

**Author notes:** Corresponding author, e-mail address, postal address: Department of Evolutionary Biology and Environmental Studies, University of Zurich, Winterthurerstrasse 190, 8057 Zurich.

## Abstract

Species are seen as the fundamental unit of biotic diversity, and thus their delimitation is crucial for defining measures for diversity assessments and studying evolution. Differences between species have traditionally been associated with variation in morphology. And yet, the discovery of cryptic diversity suggests that the evolution of distinct lineages does not necessarily involve trait differences. Here, we analyze 1,684,987 variant sites and over 4000 genes for more than 400 samples to show how a tropical montane plant lineage (*Geonoma undata* species complex) is composed of numerous unrecognized genetic groups that are not morphologically distinct. We find that 11 to 14 clades do not correspond to the three currently recognized species. Most clades are genetically independent and geographic distance and topography are the most important factors determining this genetic divergence. This lineage does not match the model of an adaptive radiation, but instead, constitutes the first example of a hyper-cryptic plant radiation in tropical mountains.

## INTRODUCTION

Understanding the characteristics of species formation is central to our interpretation of current biodiversity patterns and future predictions (De Aguiar et al. 2009; Butlin et al. 2009). Species diverge by various processes, including geographical isolation between populations or vicariance (Grant 1981; Mayr 1942, 1970, Turelli et al. 2001), ecological specialization (Coyne and Orr 2004; Arnold 2015; Nosil 2012), hybridization, and polyploidization (e.g., Rieseberg and Willis 2007; Shimizu 2022). Because species distributions often span wide ecological and geographical ranges, understanding the tempo and mode of species persistence in space and time amidst such heterogeneity is crucial to understanding what a species itself represents. So far, however, our knowledge about different modes of species diversification has been limited by the paucity of high coverage genetic information needed to assess the boundaries between species.

Mountains are one of the most important arenas for species radiations, as they impose geographical barriers that promote diversification and create elevational gradients that lead to a high diversity of habitats (Hughes and Atchinson 2015; Ebersbach et al. 2017). Geographical barriers, on the one hand, can lead to dispersal limitation (Särkinen et al. 2012) and the geographic diversification of numerous lineages (e.g., Cadena et al. 2012; Londoño et al. 2014; Lagomarsino et al. 2016). Elevational gradients, in turn, open numerous opportunities for divergent selection and local adaptation. This is the case of marginal populations that initially occurred at the species upper or lower elevation range limit, and which may adapt to new climatic conditions (Angert et al. 2008). Importantly, the dynamism of mountains leads to fluctuations in both dispersal barriers and elevation zones through time (Simpson, 1974; Hooghiemstra and van der Hammen 2004). Such dynamism alters the connectivity between species and populations (Flantua et al. 2014; Sanín et al. 2022a), and therefore also the morphological and genetic disparity between those groups (e.g., Jabaily and Sytsma 2013; Vásquez et al. 2016).

Rapid and recent diversification or evolutionary radiations are an important phenomenon in the evolution of biodiversity (Losos 2010; Hughes et al. 2015). In montane systems, plant radiations result from different speciation modes, including ecological adaptation (e.g., Hughes and Atchison 2015), progenitor-derivate speciation (e.g., Vargas et al. 2020), pollinator isolation (e.g., Lagomarsino *et al*. 2016), and geographical isolation followed by local adaptation (e.g., Eaton and Ree 2013). Understanding the relative importance of each of these speciation modes remains crucial to unravel the complexity of lineage formation and species maintenance in highly biodiverse regions of the world like the tropical Andes. However, revealing speciation modes in a taxonomic group is complicated by the requirement of having a large-scale sampling that encompasses the full distribution of the group, as well as high-resolution phylogenetic data to distinguish between different populations and eventually species. Currently, only a few montane plant groups have such genome-wide data (e.g., *Costus*: Vargas et al., 2020; *Lupinus*: Contreras et al. 2018; *Pedicularis*: Eaton and Ree 2013).

Montane plant radiations, spanning a large range of phenotypic and ecological conditions, are mostly the result of speciation mediated by island-like ecological opportunities following mountain uplift (Hughes and Eastwood 2006; Hughes and Atchison 2015; Lagomarsino et al. 2016). Little is known, however, about those montane lineages that do not conform with adaptive-innovative radiation models. Evolutionary lineages that are morphologically indistinguishable from each other are known as cryptic (Bickford et al. 2007; Struck et al. 2018) and these are important because they may represent unique, overlooked evolutionary trajectories. For this reason, research on current and future biodiversity patterns must consider the study of cryptic biodiversity (Bálint et al. 2011; Fišer et al. 2018). In fact, Adams et al. (2014) warned about the potential existence of numerous “hyper-cryptic” lineages. Hyper-cryptic groups are those in which –based on the study of multiple, independent, nuclear genes– there is an increase in species-level diversity including and beyond a four-fold increase. The study of these hyper-cryptic species complexes is paramount to achieve Global Biodiversity Assessments that approximate actual species estimates.

Cryptic diversity has been most studied in animals (e.g., birds: Krabbe et al. 2020; mammals: Nicolas et al. 2012; reptiles: Brown et al. 2012; arthropods: Miller et al. 2013; molluscs: Vriejenhoeck 2009) and more recently in plants from temperate ecosystems (Arctic: Grundt et al. 2006; Europe and Temperate Asia: Theodoridis et al. 2019; North America: Jolles and Wolfe 2012; Mastretta-Yanes et al. 2018; North America and Western Europe: Medina et al. 2012; Boucher et al. 2021). Studies of cryptic tropical plants are very scarce (e.g., Gale et al. 2018; Li et al. 2023; Pillon et al. 2014) and, to our knowledge, no assessment of a cryptic plant lineage has been carried out in the tropical mountains, except for the discovery of a small clade restricted to dry inter-Andean valleys (Gagnon et al. 2015). *Geonoma* is one of the most species rich genera of American palms (Arecaceae) with 68 species and 90 subspecies currently recognized (Henderson et al. 1995; Henderson 2011).

Species delimitation within *Geonoma* has been challenging because about one fifth of the species exhibit great morphological variation that is not consistently related to geographical or environmental location (Henderson 2011). These species may be considered species complexes or polymorphic species. The Neotropical mountains are home to one such species complex, the *Geonoma undata* lineage (hereafter *undata* complex). Henderson (2011) recognised five species and 16 subspecies within the *undata* complex, distributed from southern Mexico to central Bolivia. *Geonoma* originated ca. 18.5 (11.9–19.5) Ma, and the *undata* complex originated ca. 12.5 Ma with a crown age of about 5.3 Ma (3.8–9.2) (Roncal 2011). Unlike other *Geonoma*, the *undata* complex occupies habitats at elevations at 800– 3400 m, and previous studies inferred that this clade originated from lowland lineages through adaptation to cold environments (Roncal et al. 2005; 2011).

The lack of genomic data and sampling at the species or subspecies level has precluded our understanding of how much of the diversification of the *undata* complex is the result of adaptation or drift due to geographical isolation. Targeted sequencing to evaluate the phylogenetic relationships of the whole Geonomateae tribe showed that all species within *undata* complex clade formed polyphyletic groups not corresponding to the latest taxonomic treatment (Loiseau et al. 2019). This phylogenetic reconstruction suggested lack of genetic structure, leading to few consistent clades each covering a large geographical region. However, another study using the same target sequencing approach but focused on populations of the *undata* complex in Colombia, identified several genetic clusters, some of them even with sympatric distribution, suggesting reproductive isolation and hence speciation (Sanín et al. 2022b). Thus, the question remains as to how many genetic groups and ultimately species make up the *undata* complex throughout its wide continental distribution.

Here, we aim to understand the genetic structure of a montane lineage that might well correspond to an evolutionary radiation but whose phenotypic variation does not match that of adaptive-innovative radiations. Specifically, we tackle this by assessing the diversification of the *undata* complex by conducting a regional analysis that encompasses most of its distribution across the Neotropics. We do this by analysing a very large genomic dataset in any species complex of tropical montane plants. This dataset consists of 422 samples from 70 different locations (spanning a gradient of 30° of latitude and 2400 m in elevation) of the three more widespread species within the *undata* complex – *G. lehmannii*, *G. orbignyana and G. undata* – that are all closely related and very difficult to distinguish morphologically from one another; and an extended whole genome target sequencing covering 1,684,987 variant sites for the population genetic analyses and over 4000 genes for the phylogeny. We specifically asked: i) what is the extent of genetic structure within the *undata* complex? ii) are genetic clusters within the *undata* complex so broadly distributed as to be considered part of a single extremely variable species? or are the same clades locally restricted? iii) What are the phylogenetic relationships between the resulting clades and their distributions?

## MATERIALS AND METHODS

### Taxon Sampling

We sampled leaf tissue from 422 adult individuals located in ca. 70 sites across three species from the *undata* complex (described in Henderson 2011 and Loiseau et al. 2019): *G. undata*, *G. orbignyana*, and *G. lehmannii*. *Geonoma undata* and *G. orbignyana* are the most variable, abundant, and widespread species of the group, whereas *G. lehmannii* occurs in small and disjunct populations from Panama to Peru. These distributions explain why our sampling includes only three samples of *G. lehmannii*, 271 of *G. undata* and 148 of *G. orbignyana*. We were unable to sample the locally endemic species *G. trigona* and *G. talamancana*. We aimed to cover the latitudinal and elevational gradients of the clade, and thus sampled populations from ca. 800 m to 3400 m in elevation. As outgroups, we used individuals from closely related groups like *G. macrostachys*, and species from 14 different palm genera (Appendix C in Supplementary Material). Overall, 98% of leaf tissues came from field collections, and 2% were obtained from herbarium and botanical garden collections.

### Laboratory and Bioinformatics Protocol

The detailed protocols used for the laboratory (DNA extraction, library preparation, target capture) and bioinformatics (read trimming, mapping and SNP calling) steps were described in previous publications (de La Harpe et al. 2019, Loiseau et al. 2019). Below, we summarize a few relevant points.

DNA was extracted using the standard instructions in the DNAeasy Plant Mini Kit (Qiagen). Library preparation was done using a KAPA LTP kit (Roche, Basel, Switzerland). Libraries were quantified with a Qubit® Fluorometer v 2.2. For target capture de La Harpe et al. (2019) developed a kit that targets 4,051 genes and 133 non-genic putatively neutral regions. The pooled target capture reactions were sequenced with an Illumina HiSeq3000 sequencer in paired-end 2 x 150 bp mode.

Sequencing adapters were removed and reads were trimmed using CONDETRI V2.2 (Smeds and Künstner 2011) selecting 20 as the high-quality threshold. We used BOWTIE2 v2.2.5 (Langmead and Salzberg 2012) to map reads to the *G. undata* pseudo reference genome (NCBI project: PRJNA482221). Only reads mapping at a unique location in the genome were kept for analyses. PCR duplicates were masked using PICARD TOOLS v1.119 (). GATK v3.8 (McKenna et al. 2010) was used to base-recalibrate and realign reads around indels. SNPs were called for target regions and their surrounding 1,000 bp using UNIFIEDGENOTYPER also from GATK v3.8.

Sites were further filtered with VCFtools v0.1.16 using the options: no indels allowed, minimum quality score of 30, minimum mean depth at 10× and maximum mean depth at 100× per site, and a maximum of 50% missing data allowed per site. This resulted in a set of 1,684,987 variant sites with an average depth of 32.14× and 13.5% missing data. The whole data set containing 445 samples including the outgroup samples were used for phylogenomic analyses, while a subset of only the 422 ingroup samples of *undata* complex were used for downstream population analyses.

### PCA and ADMIXTURE Analysis

Principal Component Analysis (PCA) – We used PLINK 1.9beta6.21 to prune variant sites with linkage disequilibrium. To do this we selected a window size of 50Kb, we used a window step size of 10 bp and we pruned any variables that show an r^2^ equal or greater than 0.1 within windows. Filtered variants were used to calculate the eigenvectors and eigenvalues and produce a PCA (Fig. 1).

**Figure 1.**
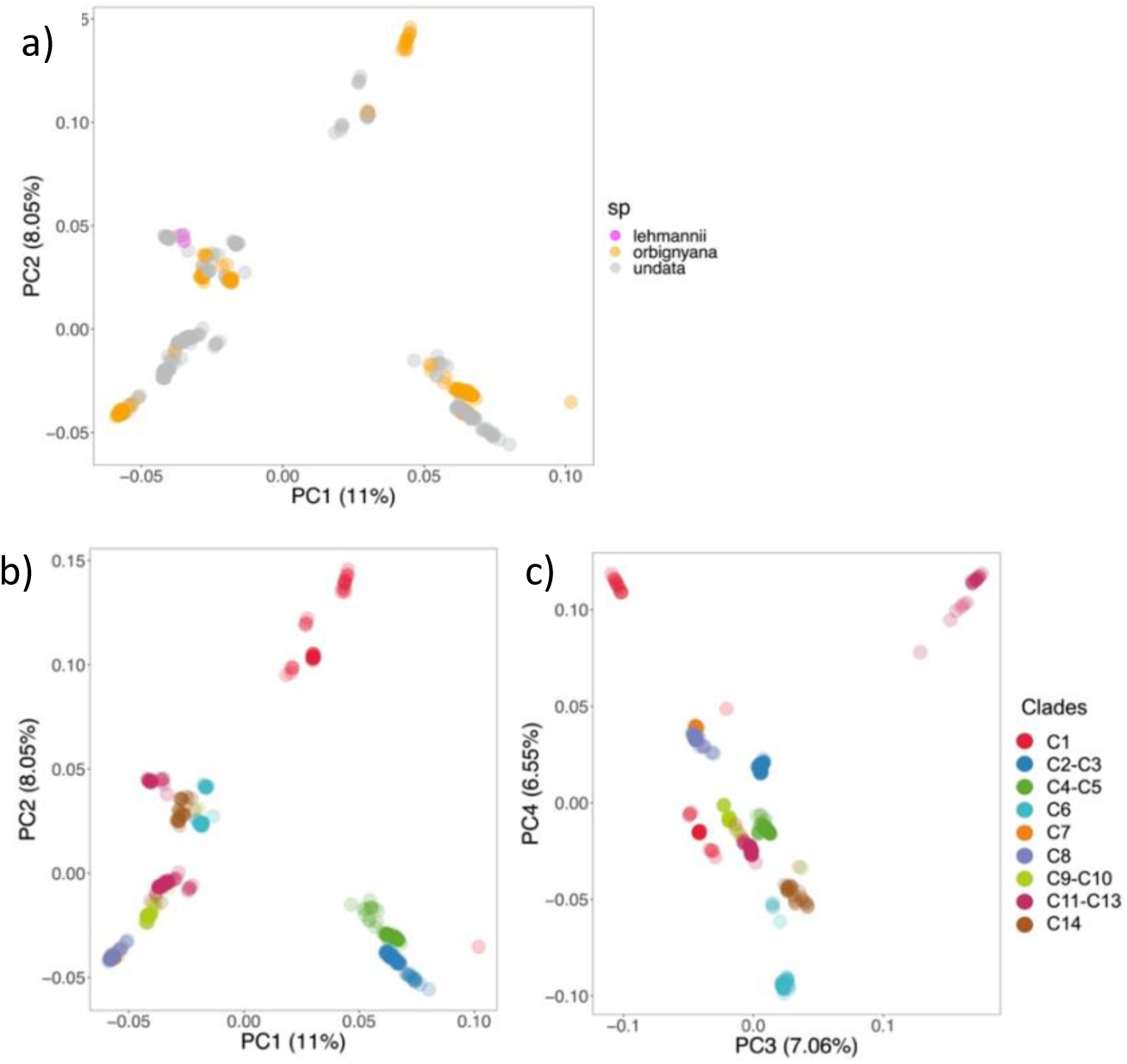
Population geonomics reveals lack of clustering in the current taxonomic classification of the *undata* complex. Principal Component Analysis (PCA) scatter diagrams show (a) populations scattered in PC1 and PC2 according to the current taxonomical classification. (b, c) Populations scattered in PC1–PC4 according to the major clades later identified in the phylogenetic analysis. Some of the clades share their color accordingly with the genetic group they belonged to in the Admixture analysis (see Fig. 2). Most of the variation along the four axes (shown in parentheses) is explained by the geographical location of each cluster (see details in the text).

Admixture – We further assessed the population structure among the *undata* complex samples using Admixture (v.1.3) (Alexander et al. 2009). We used the pruned output from the previous step which excluded the SNPs in linkage to run Admixture for a predicted number of K clusters varying between 1 and 15. We plotted the results with *Pong* (Behr et al. 2016) (Fig. 2). We used QGIS 3.4.12 to map the locations of each group.

**Figure 2.**
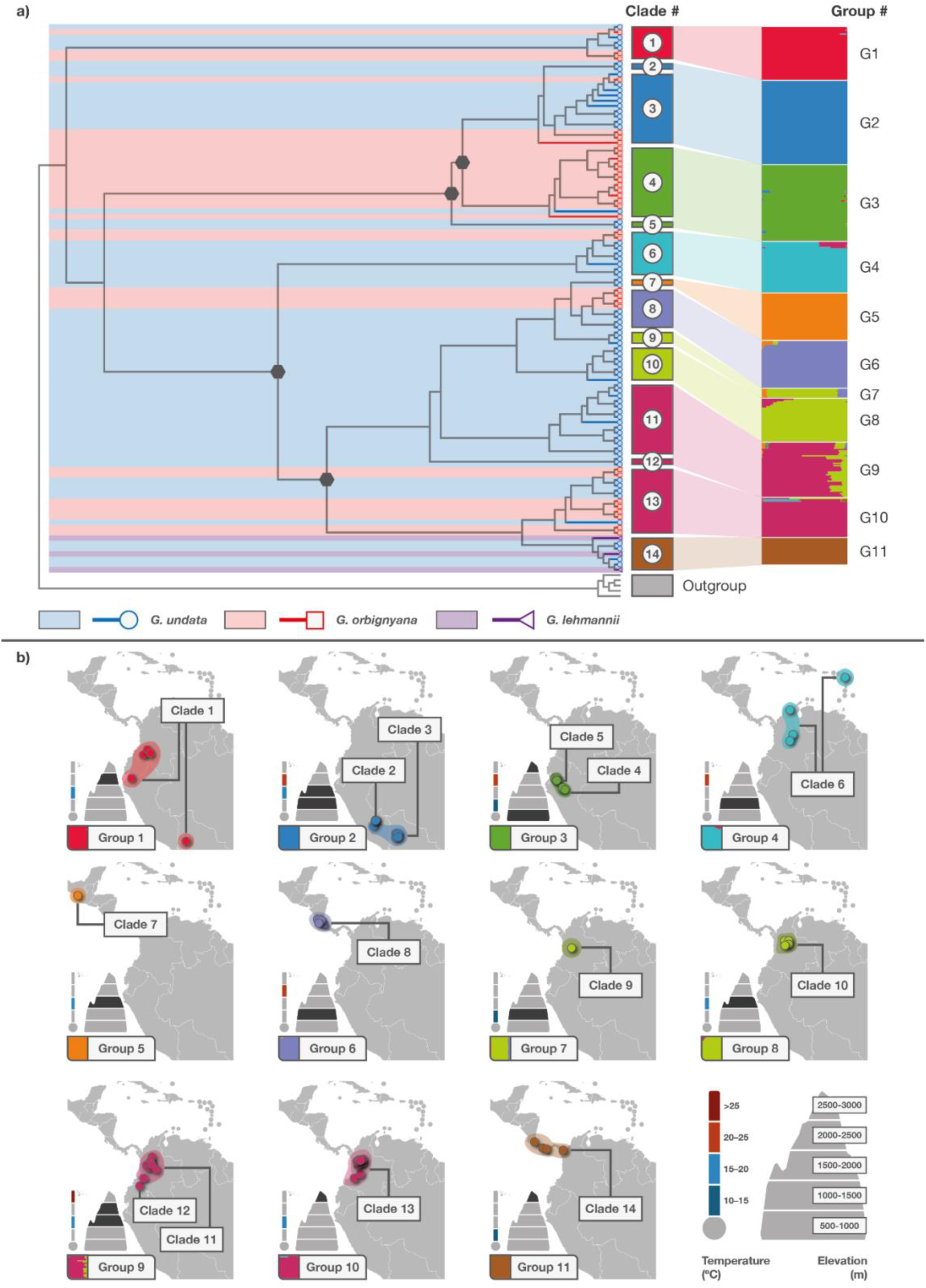
Phylogenomic and population structure analysis show that the *undata* complex consists of 11 to 14 genetic lineages with various levels of admixture and distinct geographical and elevational distributions. (a) *Left*: Simplified phylogeny including only two samples per population, tip-branches and their background are colored according to their current taxonomic classification, branches with less than 0.8 support are indicated with gray dot-hexagons. (See full tree and support values in Supplementary Material (Appendix B)).

### Divergence and Diversity Parameters

We calculated population genetic parameters including pairwise nucleotide diversity (π) (Nei and Li 1979), average number of pairwise differences (D_xy_) (Nei and Li 1979) and pairwise F_st_ (Reynolds et al. 1983). All parameters were calculated both per genetic clusters inferred from admixture analyses, as well as for geographical sampling point (populations). Relative mean genome-wide parameters were calculated from the average between genomic windows of 500 bp containing at least one variant site. Values per window were calculated using customised python scripts (from Martin et al. 2019).

### Phylogenomic Analyses

To obtain independent and equally informative genomic regions for the phylogenomic analyses, we used the following conservative approach: we split variant sites into genomic windows of 10 kb, we used these windows to produce independent trees that would later be used as “units” for a consensus tree. Only contigs with over 10 kb in length in the reference genome were considered. We found variation in the number of variant sites between windows suggesting that variant sites are not completely randomly distributed. Thus, based on the distribution of the number of variant sites per window, we selected windows across all sample sequences with an intermediate number of variants to produce the genetic trees. The reason for this is that trees with too few variant sites (genomic regions without markers) will overestimate the uncertainty in the relationship between groups, in particular the relationship between terminal branches/clades. Trees with too many variant sites will have the opposite effect.

From an initial number of 4718 windows containing SNPs across all samples, we obtained 2624 windows with 200 – 750 SNPs per window. Variant sites per window were used to produce local alignments by substituting variants sites into the refence genome sequence per samples. This was performed using vcf2phylip.py (Available at https://github.com/edgardomortiz/vcf2phylip). A maximum likelihood tree was produced for each alignment, using IQ-Tree. For each tree the option MFP to search for the best suited model and the ultrafast bootstrap option was used. Finally, the 2624 trees were used in ASTRAL (V. 5.7.8.) (Mirarab et al. 2014; Mirarab and Warnow 2015) to obtain a consensus tree and posterior-distribution support values.

### Environmental Variation

We obtained the values for the mean annual air temperature from the CHELSA V2.1 database (Karger et al. 2017) for each sample and used them to analyze the variance (ANOVA) in temperature between phylogenetic clades. We did the same analysis to test for the differences in elevation between clades.

## RESULTS

### Genetic Structure

Our Principal Component Analysis (PCA) showed that the current taxonomical classification of our samples into the species *G. lehmannii*, *G. orbignyana*, and *G. undata* is not supported by the genomic differentiation between populations (Fig.1a). Therefore, to further understand how individuals and populations cluster and relate to each other, we used Admixture and phylogenomic analyses (see results in the following sections). The clades obtained from the phylogeny were then used to plot the PCA again (Fig. 1b). We found that PC1 explained 11% of the genetic variance. Except for one population from Colombia, PC1 separated populations in a north–south gradient. PC2 explained a further 8% of the variance and separated Clade 1 (C1) (that combines populations from southern Colombia, southern Ecuador, and northern Bolivia) from the rest. PC3 and PC4 together explained 14 % of the variance and distinguished populations that occur in some of the extremes of the geographical boundaries of our sampling: the Darien region between Colombia and Panama, Tapantí National Park in Costa Rica, La Paz province in northern Bolivia, and the Sierra Nevada de Santa Marta in Colombia (Fig. 3a).

**Figure 3.**
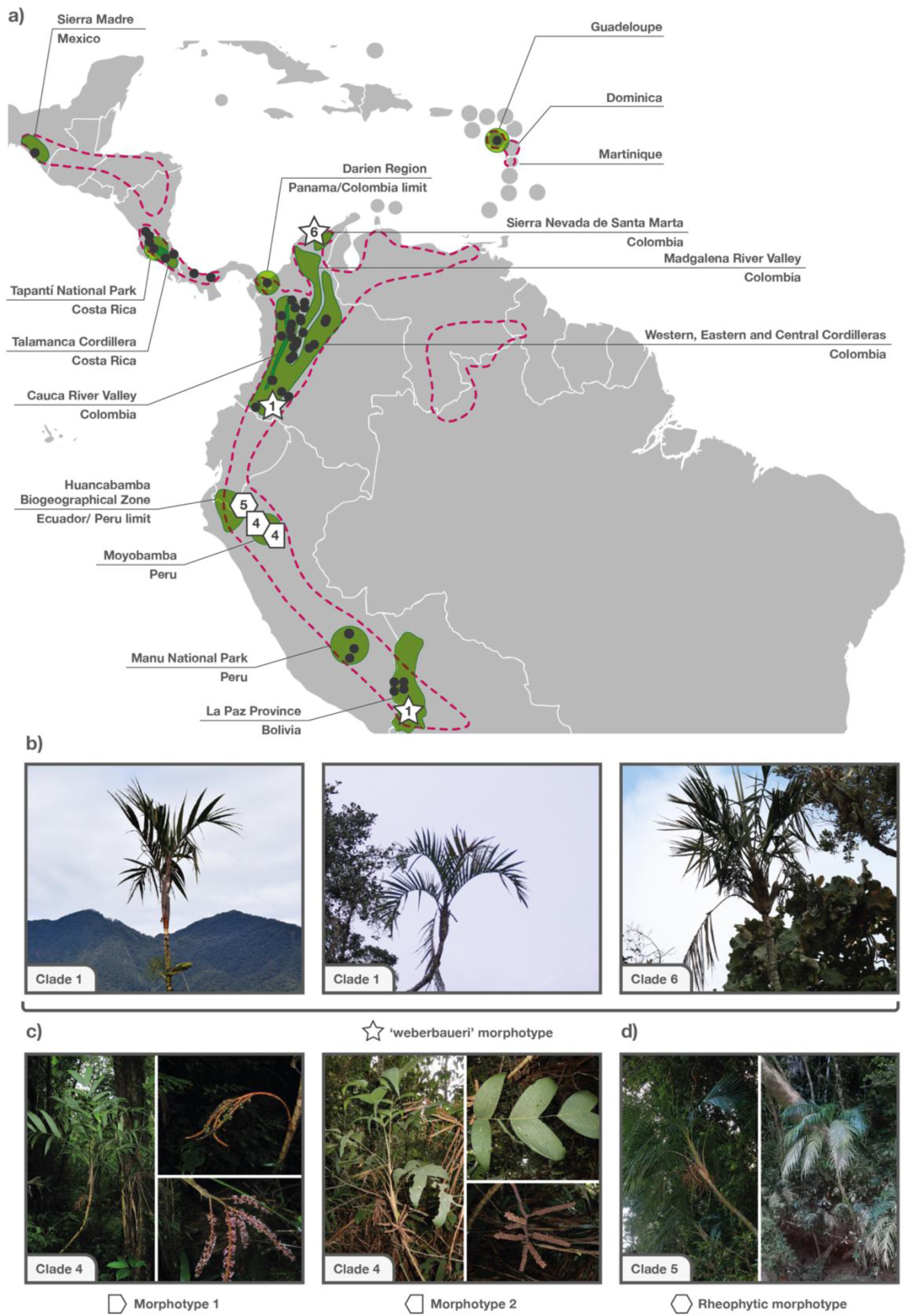
a) Geographical distribution of the *undata* complex populations sampled in this study. Location and names of specific places and topographical features mentioned in the text are indicated. Dotted-red areas indicate the whole distribution of the *undata* complex based on botanical records (Henderson, 2011). **Although there is phenotypic convergence and differences among some of the clades, our results showed that these differences might not necessarily confer evolutionary distinctiveness**. b) Morphological convergence of populations from high elevations but located in distant places. c) Differentiation in leaf division and inflorescence thickness that can be typically observed in nearby populations of the *undata* complex. d) Distinct morphology of a rheophythic population from the eastern Ecuadorian lowlands. *Right*: Admixture results showing the resultant genetic groups. Notice that although nine groups were supported by the loglikelihood values (Fig. S2 in Supplementary Material, Appendix A), these are numbered G1–G11 because we differentiated two groups (G7, G9) that had a high level of admixture as we further detail in the text. (b) Maps for the distribution of the clades and corresponding genetic groups. Mountain and temperature icons show the elevation and temperature ranges occupied by each clade.

To further understand the population structure of our *undata* sampling we examined the outcome from the Admixture analyses. We found that nine groups (K=9) had loglikelihood values with minor changes. These correspond to seven independent groups and two with high levels of admixture (Fig. 2a). Regarding the two groups with high admixture, G7 shared variation with G5, G6, and G8, whereas G9 shared variation with G8 and G10 (Fig 2a).

### Phylogenomic Structure

To understand the cluster patterns within these groups and the relationships between them, we next assessed the phylogenomic results. Based on our maximum likelihood consensus tree (Fig. 2a), we found 14 well-supported clades in the *undata* complex, although the relationships between some of the clades were not always well-supported. Our phylogeny has three basal and strongly supported splits: the first split corresponds to Clade 1 that clusters populations from distant geographical areas; the second split clusters populations from Ecuador, Peru, and Bolivia; while the third integrates populations from Colombia, the Caribbean, Central and North America. The limit between the last two clades occurs near the equator. These large clades are furthered divided into smaller clades, some of which contain populations that are geographically narrow, and others in which relatively distant populations cluster together. Below, we provide details of the geographic distribution and populations that integrate each of the 14 clades.

Clade 1: populations in three geographically remote high elevation sites, including specimens morphologically identified as both *G. orbigyana* and *G. undata*. The most divergent population is from northern Bolivia, as sister to populations from southern Ecuador and southern Colombia. We confirmed the robustness of this clade by calculating the divergence and diversity parameters (D_xy_, F_st_ and π) for the populations that compose Clade 1 and those populations that occur nearby but that clustered in different clades as this one. All parameters supported the integrity of Clade 1 by showing less genetic distance between populations within the clade than with populations from other geographically close populations (Fig S4. Supplementary Material, Appendix A).

Clade 2: a single a population of *G. undata* morphology at low elevation (ca. 1000 m) in a dwarf forest that emerges as a rather isolated mountain in the lowlands of the Peruvian Amazon in Manu National Park. Despite this clade being supported in the phylogeny as being independent, the Admixture analyses show that it is genetically related to Clade 3.

Clade 3: populations from intermediate elevations (1100–2100 m) in relatively dry habitats from southern Peru and northern Bolivia, including plants morphologically corresponding both to *G. orbigyana* and *G. undata*.

Clade 4: samples from southern Ecuador and northern Peru within the Huancabamba biogeographical zone. Most specimens in this group occur at high elevations, but it also includes a few samples from localities with special habitats and morphologies, such as a sample from a relatively isolated low peak (1498 masl) in Moyobamba, Peru. It includes plants morphologically corresponding both to *G. orbigyana* and *G. undata*.

Clade 5: a rheophytic population from southern Ecuador, corresponding to *G. undata* subspecies *pulcherrima* of Henderson (2011). In the Admixture analyses, this population is not separated from Clade 4, whereas in the phylogenetic analysis, Clade 5 is resolved as sister to Clades 1–4. The relationships of this rheophytic population are thus unclear.

Clade 6: populations from the eastern cordillera of Colombia plus populations from the Sierra Nevada de Santa Marta and one population from the Caribbean Island of Guadeloupe (= *G. undata* subsp. dussiana of Henderson (2011)). Sampled specimens in this clade included significantly different morphotypes that could all be assigned to *G. undata*, and a few also corresponded to *G. orbignyana*.

Clade 7: one medium-elevation population in southern Mexico, corresponding to *G. undata* subsp. edulis of Henderson (2011).

Clade 8: low-elevation (1000–1400 m) populations from northern Costa Rica. One population is geographically very close to another Costa Rican population (placed in Clade 14) that is present at higher elevations (>2500 m) but these two populations are significantly divergent. Morphologically, this clade includes both *G. orbignyana* and *G. undata*.

Clade 9: low-elevation samples from the western flank of the western cordillera of Colombia, morphologically corresponding to *G. undata* subsp. *stenothyrsa* (Henderson 2011). Despite geographical proximity and morphological similarity, this clade was not recovered as sister to Clade 10, but rather as sister to Clades 7 and 8 from Mesoamerica.

Clade 10: medium-elevation populations from the eastern slopes of the northern part of the central cordillera of Colombia. With only one exception, all specimens morphologically corresponded to *G. undata*.

Clade 11: medium-to high-elevation populations of *undata* morphology from the central and western Cordilleras of Colombia, from localities bordering the Cauca River Valley. This clade corresponds to the group identified by Sanín et al. (2022b) as ‘westcentral 1’, one of the sympatrically divergent groups of the *undata* complex in Colombia.

Clade 12: one high-elevation population of *undata* morphology from southwestern Colombia that is geographically not far from other population from southern Colombia assigned to Clade 1. These two populations occur, however, on opposite slopes of the Andes, and Clade 12 was recovered as sister to Clades 7–11.

Clade 13: mostly high-elevation populations from the northern part of the western and central cordilleras of Colombia. Its populations are located towards the internal slopes of the Cauca River Valley, as populations in Clade 11. It corresponds to the group ‘westcentral 2’ identified by Sanín et al. (2022b) as sympatrically divergent from Clade 11. The relationship between this Clade and the cluster conformed by Clades 11 and 12 is still to be confirmed, because in our phylogeny the support between these branches was low. In addition, Admixture analysis showed that populations in Clades 11, 12 and 13 probably experience active gene flow. In accordance with this, morphologically this clade freely mixes individuals assigned to *G. orbignana* and *G. undata*.

Clade 14: Two high-elevation groups coincide here, a low-lying group formed by samples from the lowlands (ca. 900-1400) from northern Panama to the Panama-Colombia border region, and a group including samples from the highlands of Costa Rica. Some of these samples were originally identified as *G. lehmannii*, others as *G. undata*.

In short, our phylogeny demonstrated that the *undata* complex is composed of several genetically independent groups that could be in close geographical proximity. Three major clusters were strongly supported, one with a disjunct distribution and another two encompassing a north-south gradient. These clusters were further divided into smaller clades occupying relatively small geographical areas or spread along elevation gradients, as described in the following section.

### Divergence and Diversity Parameters

The parameters D_xy_, F_st_ and π confirmed the results obtained previously with the Admixture analysis and later confirmed with the phylogeny. In general, all indexes (π, D_xy_ and F_st_) showed moderate values for all clades. Clade 1 shows the largest divergence from all other clades (Fig. S3). The largest genetic distance (D_xy_) occurs between northern and southern clades. The lowest genetic diversity (π) occurred in the southern Mexico clade (Clade 7) (Table S1).

We also calculated D_xy_ and F_st_ to test for the robustness of Clades 1, 8 and 14. We calculated these parameters for i) the populations that compose Clade 1 and populations in Clades 3, 4 and 12 which occur in sites nearby Clade 1; ii) populations in Clades 14 and 8 which also have some range overlap. All parameters supported the integrity of Clade 1 (Fig. S4), Clade 8 and Clade 14 (Fig. S5) by showing that the relationship between the populations that comprise each clade is stronger than with the other geographically close populations.

### Environmental Variation

Some of the genetic groups identified by our previous analysis of the *undata* complex occupied habitats with contrasting temperatures due to their non-overlapping distribution (isolation) along elevational gradients. The analysis of variance showed that there are significant differences in the elevation (R^2^=0.5757; F=39.45; p-value < 0.005) and temperature (R^2^=0.6014; F=43.87; p-value < 0.005) values of the sites occupied by the different clades. While some clades occur in the warm lowlands (hereafter lowlands refer to <1500 masl), others mostly occur in the cooler highlands (hereafter highlands refer to >1500 masl) (Fig. 2b). In some cases, these clades showed little population structure and differentiation. This was the case for Clade 2 (a lowland population) found in the same genetic cluster with Clade 3 (occurring at higher elevations) and that of Clade 5 (a rheophytic lowland population) clustering with Clade 4 (Fig 2a). In other cases, however, geographically overlapping, but phylogenetically more remote clades were separated by elevation. This was the case, e.g., for Clades 8 and 14 in Costa Rica, and Clades 9 and 10 in Colombia. Overall, these results suggest that there might be environmental adaptation involved in the differentiation of some clades, but confirmation of this will require further investigations, as we discuss below.

## DISCUSSION

Here, we attempted to understand the genetic structure of a tropical montane lineage whose limited phenotypic variation does not match that of an adaptive-innovative radiation, where diversification is driven by morphological and ecological divergence. Our results demonstrate that the palms of the *undata* complex better correspond to a model of hyper-cryptic speciation and that, as a result, about a dozen of species might exist instead of the three species currently recognized and studied here. In other words, the morphological and geographical arguments currently used to delimit species in this complex and probably others in the genus (Henderson 2011) require a complete re-evaluation to achieve a robust classification. In the following sections, we discuss the implications of our results and highlight future research avenues that will contribute to resolve species delimitation in these and other hyper-cryptic mountain lineages.

### Genetic Structure within the Undata Complex

Our population structure analyses revealed limited gene flow between the different populations, even among those in the same mountain range. A combination of topography, dispersal limitation, and environmental conditions appear to drive the divergence between members of the *undata* complex.

In contrast to iconic examples of Andean plant radiations, like *Lupinus* (Hughes and Atchinson, 2015) and bellflowers (Lagomarsino et al. 2016) where species diversification is characterised by multiple changes in growth form, habitat preferences, and pollination syndromes, the diversification of the *undata* complex has been much more veiled. The three species included by us are traditionally differentiated only by two subtle traits: the shape and texture of the inflorescences first-bract. Our findings thus suggest that the *undata* complex matches the definition of a hyper-cryptic species complex in which, due to genetic differences, there is a four-fold or higher increase in species number that is largely not evident morphologically (Adams et al. 2014). However, two clades (Clades 2 and 5) seem to derive phylogenetically from a progenitor-derivate pattern of speciation (Crawford 2010).

Individuals in these clades occupy the lowest elevations recorded in our sampling and their morphology is the more distinctive of all our specimens. Still, neither Admixture, D_xy_, F_st_, or π supported a strong divergence or reduced diversity of Clades 2 and 5 in relation to their sister clades (Fig. S3 and Table S1 in Supplementary Material, Appendix A). Therefore, the hypothesis that phenotypic differentiation of these clades confers them evolutionary distinctiveness remains to be tested.

### Morphological Variation

Henderson (2011) highlighted the difficulty to classify the morphological variation within the *undata* complex. Based mostly on inflorescence characters, he recognized five species: *G. undata*, *G. orbignyana*, *G. lehmannii*, *G. trigona*, and *G. talamancana.* Using a combination of stem and leaf characters, he further distinguished 10 subspecies of *G. undata* and two of each *G. orbignyana* and *G. lehmannii*. Our study shows that this classification does not correspond to genomic groups, and that *G. orbignyana* and *G. undata* are widely intermixed in the genomic groups, with *G. lehmannii* placed in a phylogenetically distinct clade but intermixed with plants morphologically assigned to *G. undata*. This suggests that the morphological traits used by Henderson (2011) are probably not phylogenetically and taxonomically informative, and that a novel taxonomic classification is needed. Interestingly, some of the clades that we identified appear to correspond to subspecies recognized by Henderson (2011). For example, the distribution of Clade 5 corresponds to G. *undata* subsp*. pulcherrima* and Clade 9 to *G. undata* subsp*. stenothyrsa*, showing that these clades are morphologically distinctive (See Clade 5 pictures in Fig. 3d).

Our phylogenomic results showed that geographically close populations occurring at different elevations are often genetically isolated. This elevational isolation likely affects the adaptation of the species in the *undata* complex to the environment, which might manifest in some contrasting morphologies. For example, we found that the distinct high-elevation plants corresponding to the morphotype *weberbaueri* of *G. undata* subsp. *undata* (Henderson 2011) occur at distant geographic sites and fall into distinct genetic clades. Thus, *weberbaueri*-type plants from the Sierra Nevada de Santa Marta in northern Colombia were recovered in Clade 6, those from southern Colombia in Clade 1, and those from northern Bolivia in Clade 1 (Fig. 3b). As already suggested by Henderson (2011), this suggests convergent morphological evolution between populations as the result of adaptive selection.

As mentioned above, the limited morphological variation in the *undata* complex is not consistent across topographic or environmental gradients. Thus, while there are cases like the example above in which there seems to be convergence of high elevation populations or those in which a specific morphotype can be expected under certain environmental conditions (e.g., Clade 5: rheophytes with tall and stout stems, and leaves with numerous narrow pinnae, Fig. 3d), there are other cases in which populations exhibit little morphological variation — relative to their sister groups— other than in the size of different organs (e.g., Clade 2, a lowland population with plant size, leaf, and inflorescence division above the average within the *undata* complex). More studies will be needed to assess whether such morphological differences actually relate to species boundaries.

### Phylogenetic Relationships Between Clades and Their Distributions

In the phylogeny, the most divergent clade (Clade 1) has a peculiar distribution (with disjunct populations in Colombia, Ecuador, and Bolivia) and is sister to all other clades. Although we cannot exclude the possibility that further unsampled populations of this clade occur in between our records, the fact that numerous populations assigned to other clades exist at geographically intermediate locations support the notion that this is a genetically distinct, but geographically disjunct clade. The distribution of Clade 1 is peculiar because, as detailed below, the rest of the clades are consistently separated in northern and southern groups. Future research that includes more populations and perhaps a whole genome sequencing approach would help to elucidate the origin of the relationship between these distant populations.

Regarding the remaining clades, the primary geographical division is between a northern and southern group. This boundary coincides with the Huancabamba biogeographical zone locates between southern Ecuador and northern Peru (ca. latitude 5° S) (Fig 3a). Several studies suggest that the distinct climatic conditions in this area, characterized by extreme precipitation gradients, strong winds, and an atypically low treeline elevation, act as a dispersal barrier for montane plants (Simpson 1975; Ayers 1999; Cosacov et al. 2009; Jabaily and Sytsma 2013; Vargas et al. 2017, Contreras et al. 2018) and animals (Bonaccorso 2009; Chaves & Smith 2011; Gutiérrez-Pinto et al. 2012). But it has also been demonstrated that the area offers a diversity of habitats that contribute to the diversification of several plant groups (Weigend 2002). We found clades to the north (Clades 6–14) and south of the Huancabamba zone (Clades 2–3) as well as clades that overlap its range (Clades 4–5). Our findings suggest that the diversification of the *undata* complex has been significantly influenced by the Huancabamba biogeographical zone, which probably acted both as a dispersal barrier and as an arena for species divergence.

Further subdivisions resulted in the recognition of several major genetic clusters largely found in geographically distinct regions, and often corresponding to separate mountain regions. For instance, southern clades occur on the limits between Ecuador and Peru (Clades 4–5) and between Peru and Bolivia (Clades 2–3), respectively, whereas the northern clades correspond to a Mexican clade that occurs in the southern Sierra Madre (Clade 7), a clade from the Talamanca Cordillera in Costa Rica (Clade 8), one that extends from Costa Rica to northwestern Colombia (Clade 14). These groups thus correspond to geographically isolated populations in distinct mountain ranges, suggesting a role of geographical (allopatric) speciation.

In the Colombian Andes, the genetic structure is more complex. Here, three parallel mountain ranges are inhabited by six clades. Clade 6 groups the populations from Colombia’s eastern cordillera and the Caribbean and does not overlap with other clades. That a similar pattern exists in *Lupinus* (Contreras et al. 2018) and other taxa (e.g., Cadena et al. 2007; Gutiérrez-Pinto et al. 2012) suggests strong isolation and dispersal limitation between the central and eastern cordilleras separated by the Magdalena River valley. The other five clades (9–13) occupy Colombia’s western and central cordilleras and appear to mostly have relatively small geographical ranges. This pattern of isolation at small scales has also been found in other montane palm lineages in Colombia (Sanín et al. 2022c) and is likely due to the intricate topographic history of Colombia’s cordilleras. We cannot exclude the possibility that sampling of more intermediate populations would reveal a gradient of hybrid zones between these apparently isolated groups. Still, there is evidence that the clades found here have some degree of evolutionary independence even if they are geographically close, which is likely the result of reduced gene flow caused by the dissected topographical conditions of the Andean mountains.

### Species Delimitation

We propose that the *undata* complex consists of several clades that are genetically so distinct even in close geographical proximity to other clades (e.g., Clades 9 to 13) that they are likely independent. However, there is also evidence of gene flow between some of the clades, suggesting that species boundaries are not yet fully formed. We suggest that the resulting clades likely correspond to roughly one dozen biological species. Considering that our sampling did not include some geographically highly restricted and morphologically distinct populations recognized as subspecies by Henderson (2011), the actual number of species in the *undata* complex may even be higher.

We refrain from proposing a new classification of the *G. undata* group for three reasons. First, although our sampling was extensive, we did not include samples from all subspecies recognized by Henderson (2011). Second, our results do not provide direct evidence for reproductive isolation of the clades, which would be a major criterion for species delimitation. In particular, the independence of spatially proximate populations, especially those at different elevations on a single mountain chain, remains to be elucidated. Third, the genomic groups that we identified here should be evaluated from a phenotypic perspective, with the hope of detecting traits whose taxonomic relevance has so far not been appreciated. Singhal et al. (2018) in their *framework to resolving cryptic species* provided four steps to diagnose species boundaries across cryptic lineages: 1) statistical species delimitation, 2) *post hoc* discovery of phenotypic differences between clades, 3) indirect or direct estimates of evolutionary isolation between clades, and 4) calibration-based approaches. The present study contributes to step 1 and can be considered as the first step to determine which lineages are unique at the regional scale. To tackle step 2, future studies will have to assess morphological or other phenotypic (e.g., physiological) characters that might relate to species distinctiveness. Step 3 will require future research of genetic structure and reproductive biology across different elevational or climatic gradients. Finally, calibration approaches (step 4) combine information on genetics, morphology, and reproductive isolation directly from contact zones to delimit species boundaries. The latter is an appealing approach that will, however, require the compilation of considerable information and which could constitute a long-term objective in the study of hyper-cryptic lineages.

## CONCLUSIONS

Our study shows the *undata* complex is an example of an active hyper-cryptic radiation. Current morphological classification of the *undata* complex was not supported by the genetic groups identified here. Instead of three species, the samples studied here might better correspond to about dozen species. Our results support a strong population structure with little gene flow between most of the genetic clusters. We found a combination of early divergent clades that might preserve the ancestral genetic background and other genetic groups that likely represent well-defined species. A few groups showed evidence of gene flow which might later lead to further species differentiation or to the unification of current incipient species. We propose that topographical features, dispersal limitation, and environmental changes along elevational gradients are the main factors driving the diversification of the *undata* species complex.

## ACKNOWLEDGEMENTS

We thank Oriane Loiseau, Anna Weigand, Marylaure de la Harpe, Jonathan Rolland and all collaborators of the POPCORN project in Colombia, Ecuador, Peru, and Bolivia for assistance during fieldwork, permit application, and lab work, Henrik Balslev, Rodrigo Bernal, Angela Cano, and Gloria Galeano for sharing samples, and especially Margot Paris for designing and performing the targeted sequencing analyses, and for the preparation of the genetic dataset.

## APPENDIX A

### Genetic structure analyses

The first split in our admixture analyses distinguishes the northern samples from southern samples where the limit between groups is around the equator. With 15 clusters (K=15) Admixture showed one division for each sampled population (Fig S1). The loglikelihood plot showed smaller changes occur at nine clusters (K=9) (Fig S2).

**Figure S1.**
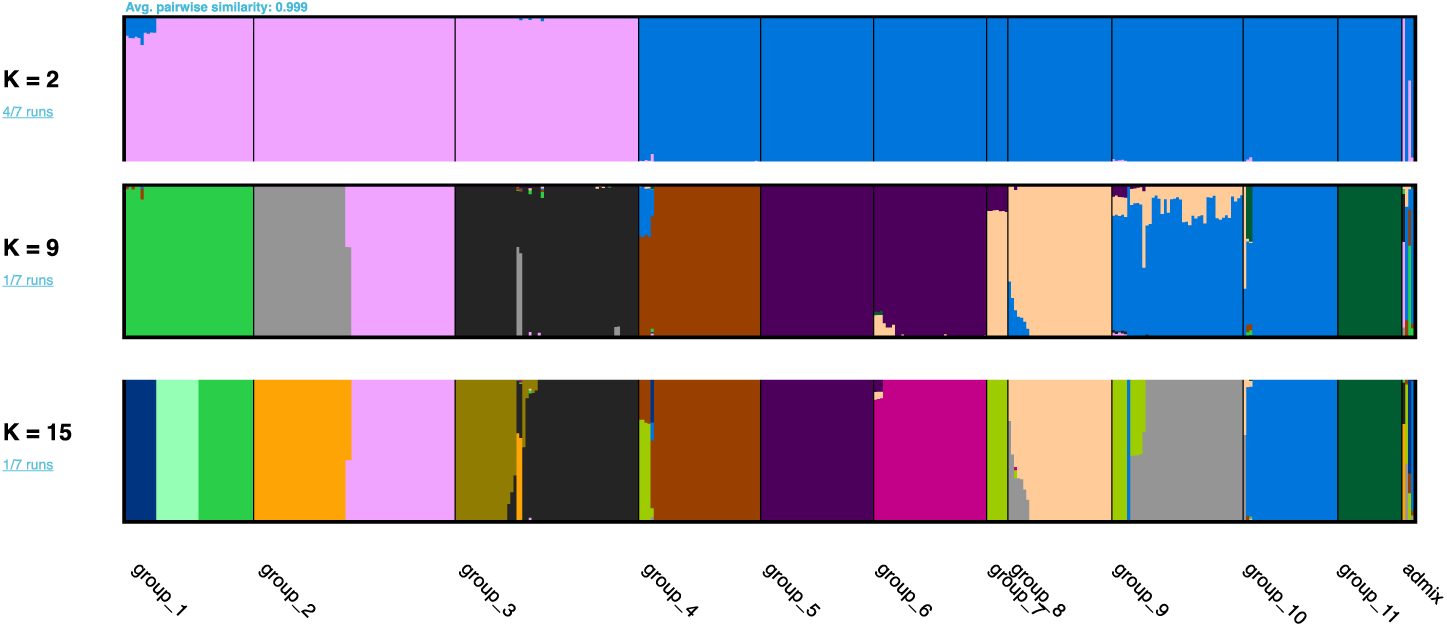
Genetic structure of sampled populations of the *undata* complex for clusters 2 to 15.

**Figure S2.**
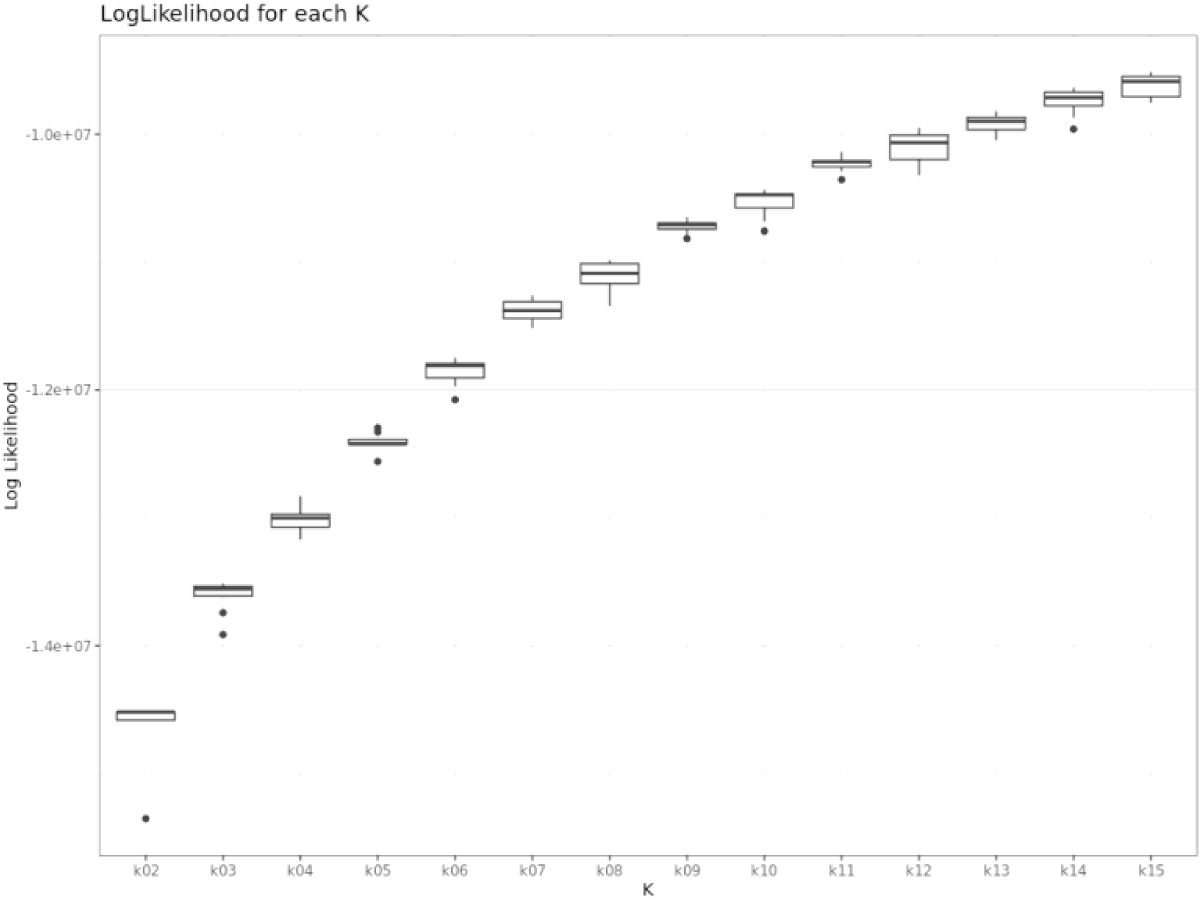
Loglikelihood boxplots for the ADMIXTURE groups from K2 to a max of K15.

**Figure S3.**
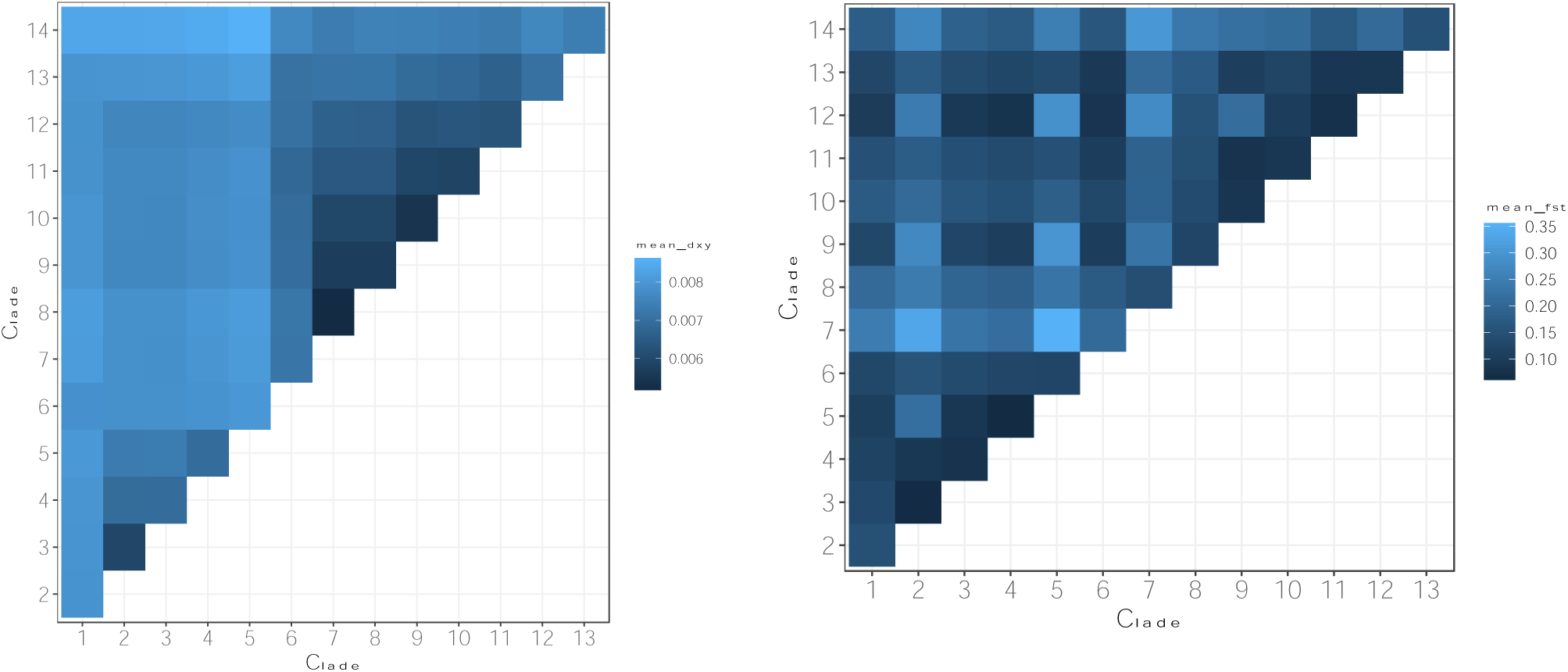
Heatmaps of the divergence measurements Dxy (a) and Fst (b) for all clades obtained in the phylogenomic analyses.

**Figure S4.**
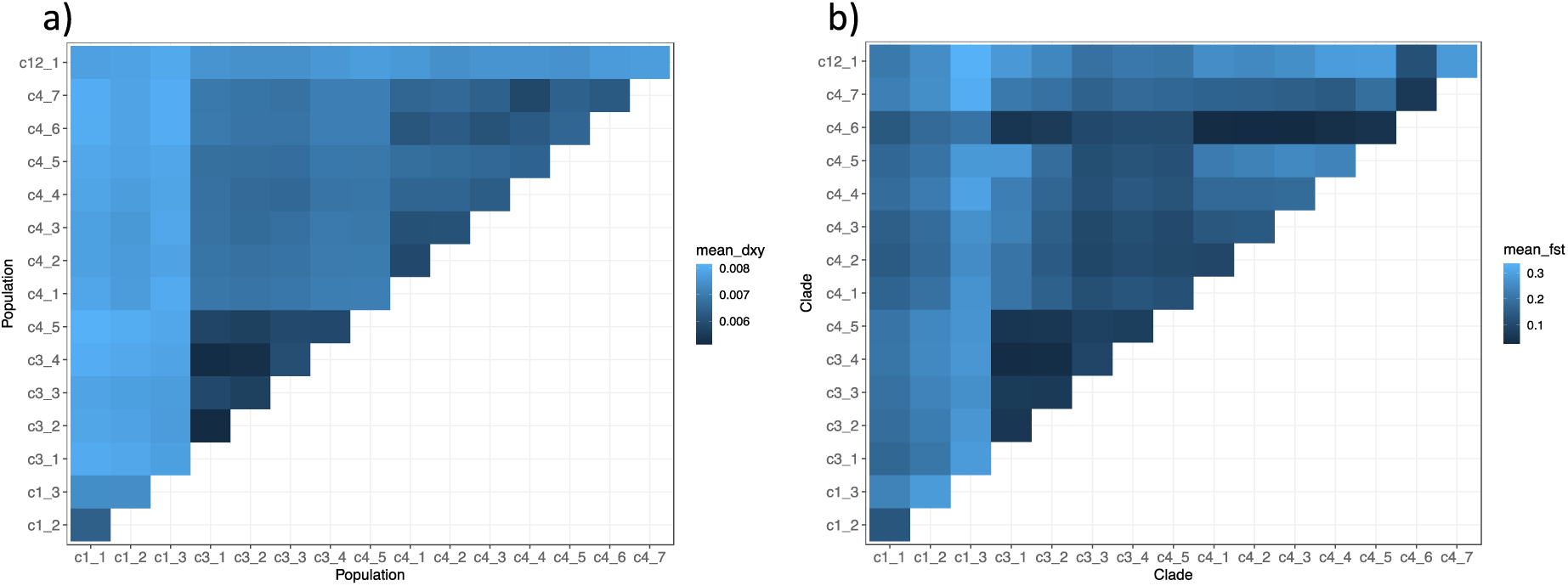
Heatmaps of the divergence measurements Dxy (a) and Fst (b) for the populations that compose Clades 1, 3, 4 and 12 which occur in geographical nearby areas.

**Figure S5.**
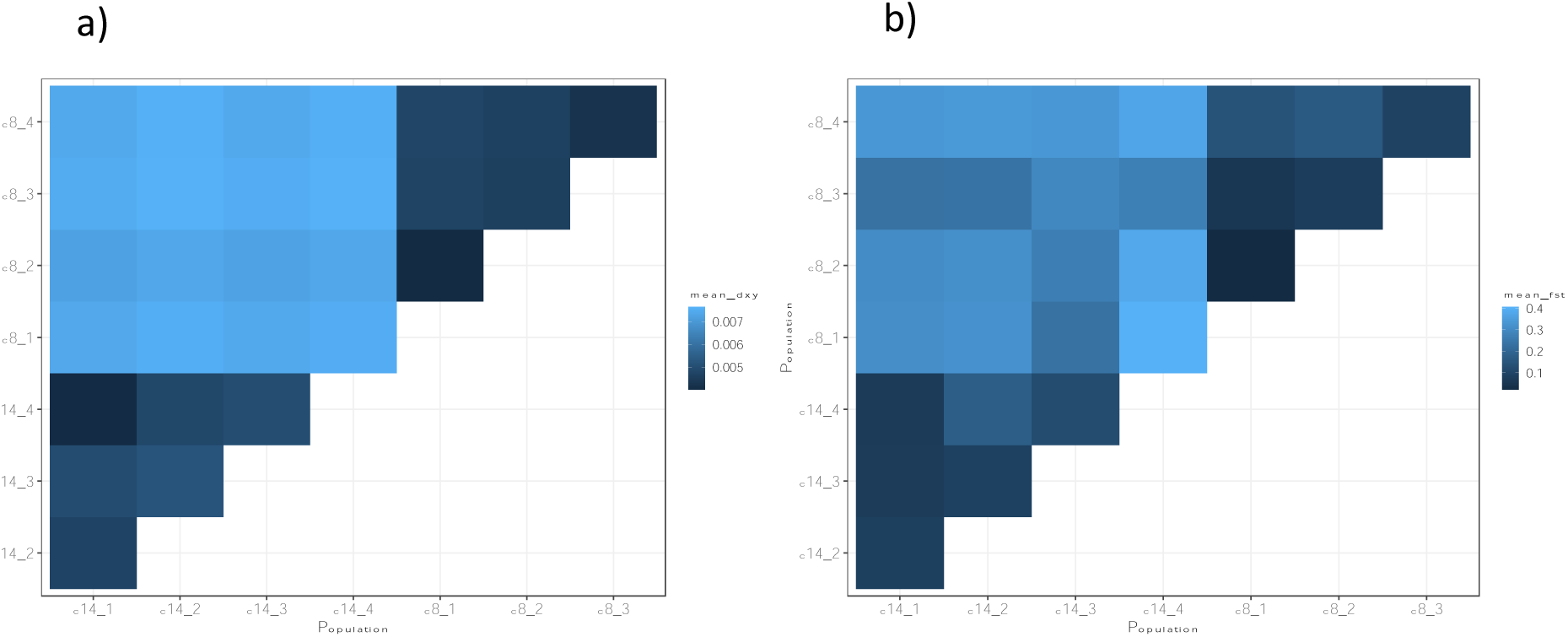
Heatmaps of the divergence measurements Dxy (a) and Fst (b) for the populations that compose Clades 8 and 14 which occur in geographical nearby areas. *Envionmental Variation: ANOVA Analyses*

**Table S1.**
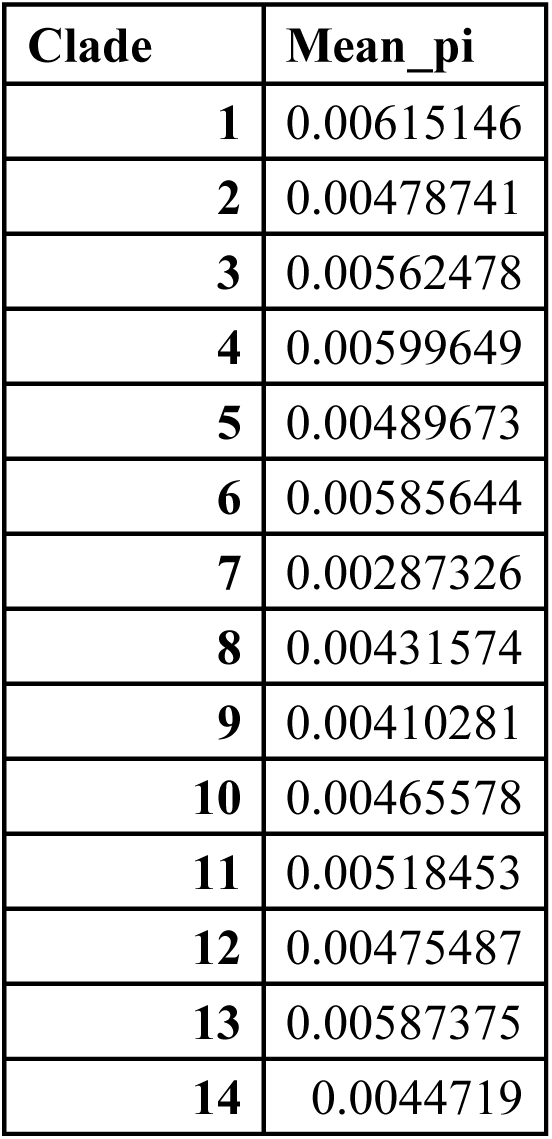
Average pi values for each of the clades supported by the phylogeny.

**Table S2.**
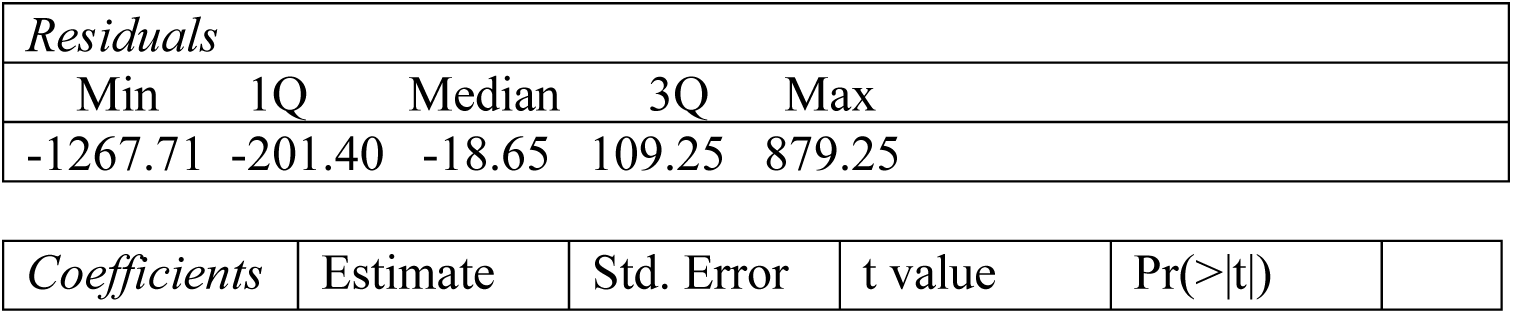
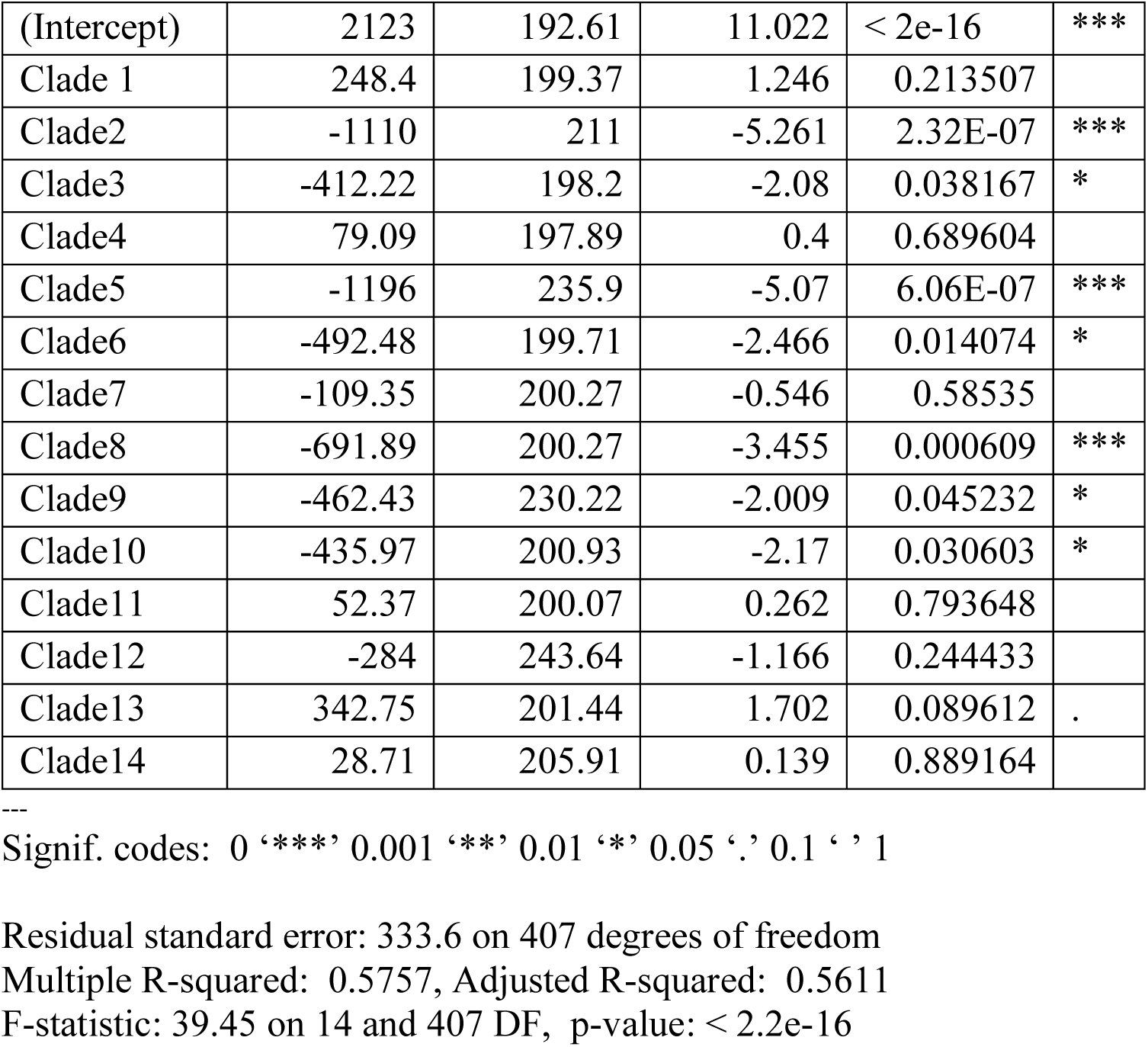
Coefficients and summary results of the ANOVA for the differences in *elevation* between clades.

**Table S3.**
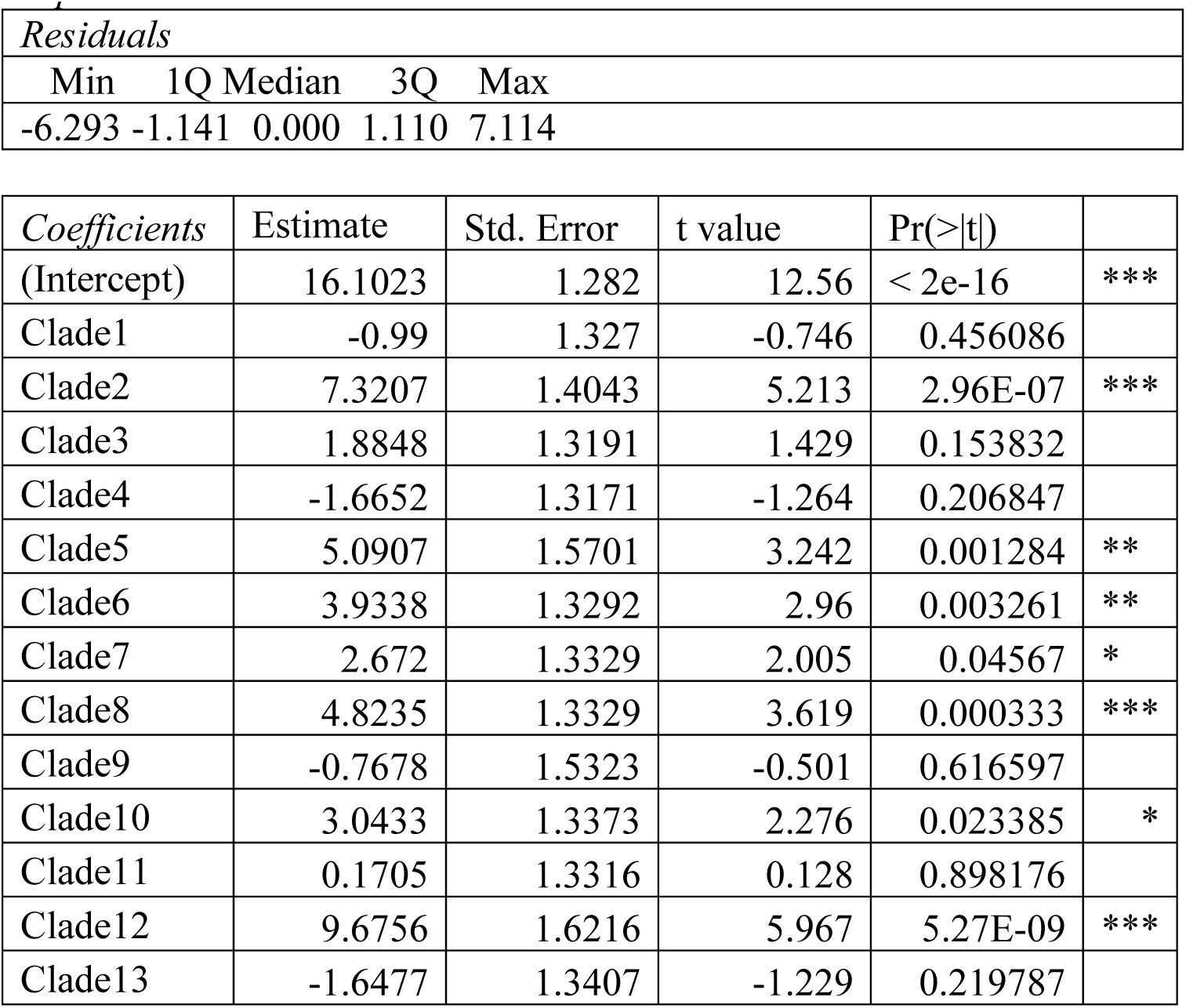
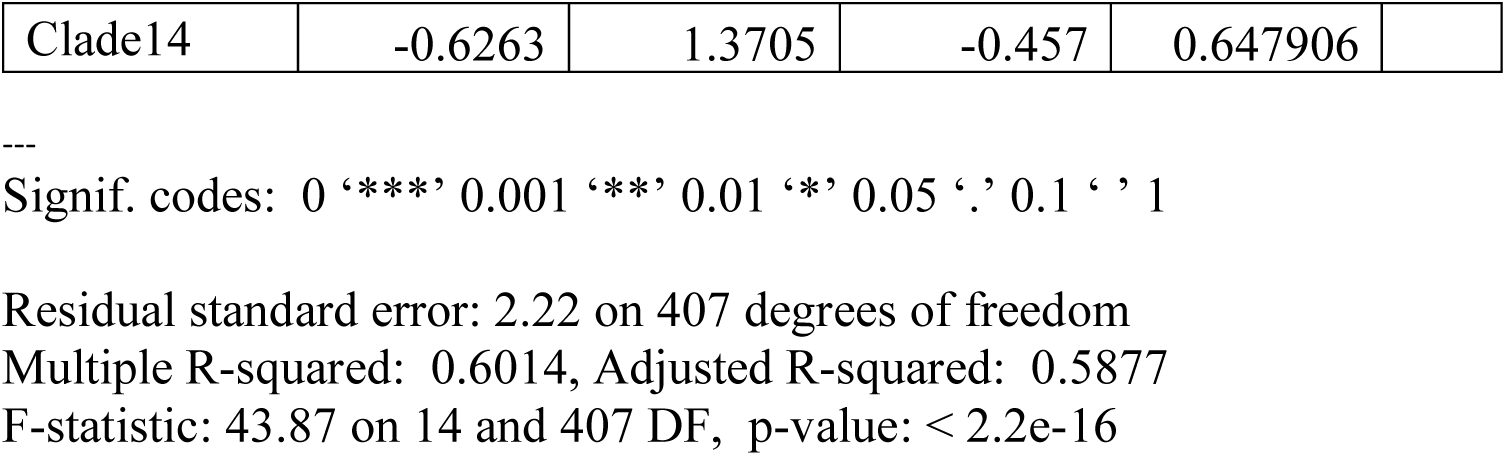
Coefficients and summary results of the ANOVA for the differences in *temperature* between clades.

## APPENDIX B

Complete phylogeny in a separate .pdf file.

## APPENDIX C

List with species names for all samples used in the phylogeny in a separate excel file.

